# Sound context modulates perceived vocal emotion

**DOI:** 10.1101/2020.01.08.898205

**Authors:** Marco Liuni, Emmanuel Ponsot, Gregory A. Bryant, JJ Aucouturier

## Abstract

Many animal vocalizations contain nonlinear acoustic phenomena as a consequence of physiological arousal. In humans, nonlinear features are processed early in the auditory system, and are used to efficiently detect alarm calls and other urgent signals. Yet, high-level emotional and semantic contextual factors likely guide the perception and evaluation of roughness features in vocal sounds. Here we examined the relationship between perceived vocal arousal and auditory context. We presented listeners with nonverbal vocalizations (yells of a single vowel) at varying levels of portrayed vocal arousal, in two musical contexts (clean guitar, distorted guitar) and one non-musical context (modulated noise). As predicted, vocalizations with higher levels of portrayed vocal arousal were judged as more negative and more emotionally aroused than the same voices produced with low vocal arousal. Moreover, both the perceived valence and emotional arousal of vocalizations were significantly affected by both musical and non-musical contexts. These results show the importance of auditory context in judging emotional arousal and valence in voices and music, and suggest that nonlinear features in music are processed similarly to communicative vocal signals.

## 1. Background

When animals are highly aroused there can be many effects on their bodies and behaviors. One important behavioral consequence of physiological arousal is the introduction of nonlinear features in the structure of vocalizations (Briefer, 2012; Fitch, Neubauer, & Herzel, 2002; Wilden et al., 1998). These acoustic correlates of arousal include deterministic chaos, subharmonics, and other non-tonal characteristics that can give vocalizations a rough, noisy sound quality. Nonlinear phenomena are effective in communicative signals because they are difficult to habituate to (Blumstein & Recapet, 2009), and they successfully penetrate noisy environments (Arnal et al. 2015).

Alarm calls and threat displays in many species often have such nonlinear features. In humans, alarm signaling manifests as screaming. Screams are characterized by particularly high signal-to-noise ratios at the most sensitive frequencies of human hearing (Begault, 2008). Recent psychophysical and imaging studies suggest that vocal sounds containing low-level spectro-temporal modulation features (i.e., modulation rate of 30-150 Hz) are perceived as rough, activate threat-response circuits such as the amygdala, and are more efficiently spatially localized (Arnal et al., 2015). But what is the effect of sound context on the perception of vocal screams? Many variable internal and external factors can potentially affect subjective judgments of a stimulus, and cues from multiple modalities can give contradictory information (Abitol, et al., 2015; Driver & Noesselt, 2008). In humans, high-level cognitive processes such as the emotional or semantic evaluation of a context can change the way signals are integrated across sensory areas (Blumstein et al., 2012; Mothes-Lasch et al., 2012). For example, musical pieces with added distortion are judged differently on the two emotional dimensions of valence and arousal (termed here *emotional* arousal) from the same pieces without the noise, but the difference in emotional arousal disappeared when presented in a benign visual context (i.e., video with very little action; Blumstein et al., 2012).

These results suggest that visual information can reduce the arousing impact of nonlinear acoustic features, potentially dampening fear reactions. But very little is known about the effect of sound context on auditory perception: such context effects may be particularly strong in music where indicators of vocal arousal likely take a different semantic or stylistic meaning than in more ecologically valid situations (Neubauer et al., 2004). Belin and Zatorre (2015) described the example of enjoying certain screams in operatic singing. In a more formal experimental setting, fans of “death metal” music reported experiencing a wide range of positive emotions including power, joy, peace, and wonder, despite the aggressive sound textures associated with this genre (Thompson, Geeves & Olsen, 2018). Other examples include the growl singing style of classic rock and jazz singers, or carnival lead voices in samba singing (Sakakibara et al. 2004).

Here we examined the relationship between manipulated cues to vocal arousal, perceived emotional valence and arousal, and the sonic context in which the vocal signal occurs. We created two musical contexts (clean guitar, distorted guitar) and a non-musical context (modulated noise), in which we integrated nonverbal vocalizations (specifically, screams of the vowel /a/) at varying levels of portrayed vocal arousal. Based on the work described above, we expected that aroused voices in a musically congruent context (distorted guitar) would be judged less emotionally negative and less emotionally arousing than voices presented alone, or presented in musically incongruent contexts (clean guitar, noise).

## Methods

### Participants

23 young adults (12 women, mean age = 20.8, *SD* = 1.3; 11 men, mean age = 25.1 *SD* = 3.2) with self-reported normal hearing participated in the experiment. All participants provided informed written consent prior to the experiment and were paid 15€ for their participation. Participants were tested at the Centre Multidisciplinaire des Sciences Comportementales Sorbonne Université-Institut Européen d’Administration des Affaires (INSEAD), and the protocol of this experiment was approved by the INSEAD Institutional Review Board.

### Stimuli

Two singers, one man and one woman, each recorded nine utterances of the vowel /a/ (normalized in duration, ∼ 1.6 s.), at three different target pitches (A3, C4 and D4 for the man, one octave higher for the woman), with three target levels of portrayed anger / vocal arousal (Figure 1 - *voice* column). The files were recorded in mono at 44.1 kHz/16-bit resolution with the Max software (Cycling ‘74, version 7).

**Figure 1:**
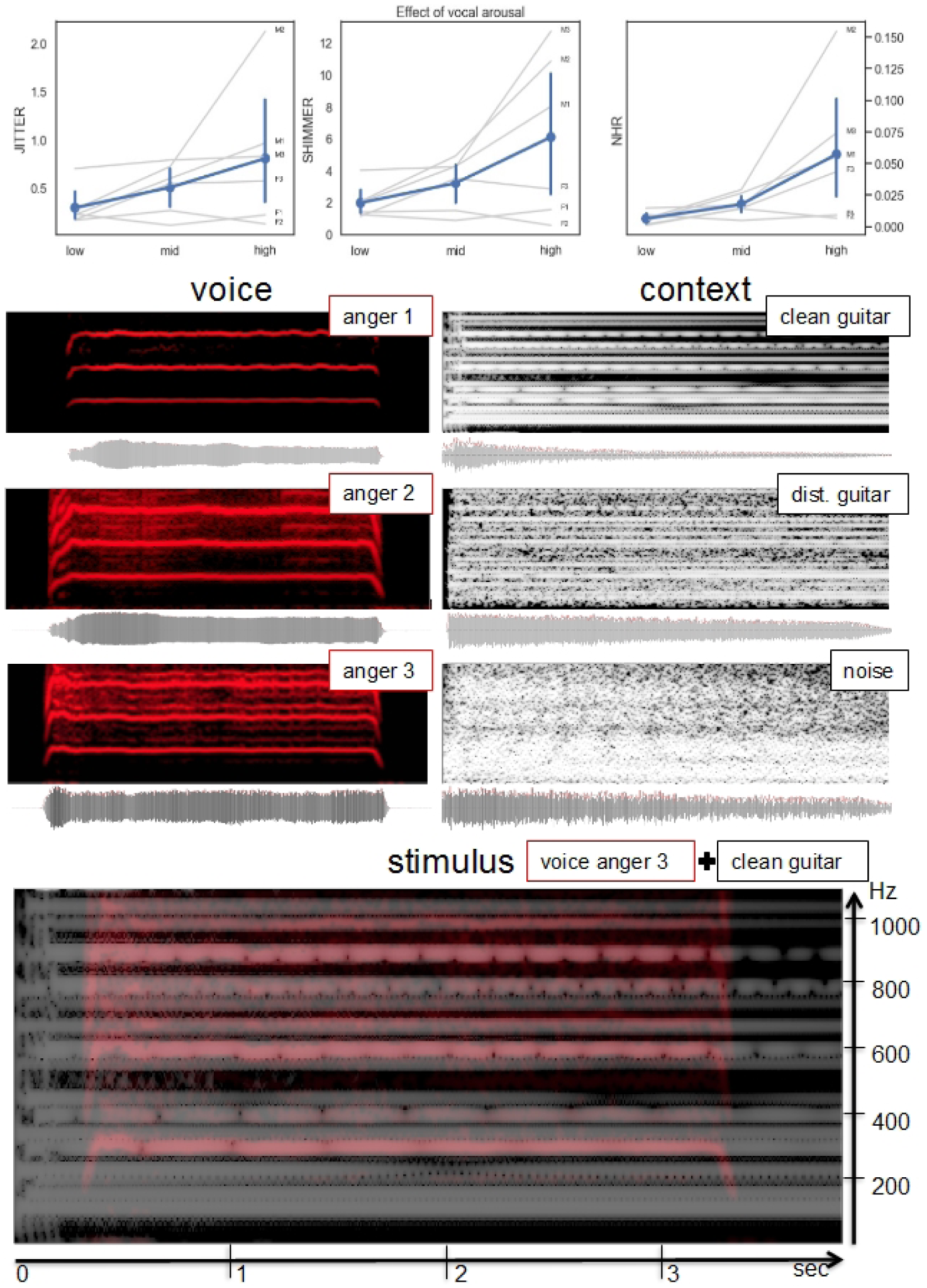
Top: Acoustical analysis (jitter_loc, vshimmer_loc and noise-harmonic ratio) of the vocalizations, as a function of level of portrayed vocal arousal. Middle: spectrograms of selected original recordings. Left column shows a male vocalization, of the note D4 at 3 anger levels. Right column shows a D-G-A guitar chord in the different sound contexts. Bottom: one example of the resulting mix.

A professional musician recorded nine chords with an electric guitar, on separate tracks (normalized in duration, ∼2.5 s.). All the chords were produced as permutations of the same pattern (tonic, perfect fourth, perfect fifth), harmonically consistent with the above vocal pitches. The nine guitar tracks were then distorted using a commercial distortion plugin (Multi by Sonic Academy) with a unique custom preset (.fxb file provided as supporting file, see the section *Data accessibility*) to create nine distorted versions of these chords. Lastly, we generated nine guitar-shaped noise samples by filtering white noise with the spectral envelope estimated frame-by-frame (frame size = 25 ms) on the distorted guitar chords (see Figure 1- *context* column).

Both vocalizations and backgrounds (noise, clean guitar, distorted guitar) were normalized in loudness using the maximum value of the long-term loudness given by the Glasberg and Moore’s model for time-varying sounds (Glasberg & Moore, 2002), implemented in Matlab in the Psysound3 toolbox. The vocalizations were normalized to 16 sones for the first level of anger, 16.8 (=16*1.05) sones for the second level, and to 17.6 (=16*1.05*1.05) for the third and highest level. All the backgrounds were set to 14 sones. For a more intuitive analysis of the sound levels reached after this normalization, we evaluated the level of the vocalizations and backgrounds composing our stimuli a posteriori. The vocalizations reached very similar levels (with differences < 2 dB among the different levels of anger), as did the different backgrounds (differences < 1.5 dB). Therefore, the overall signal-to-noise ratio (SNR) between voice and backgrounds was similar across the three different contexts (SNR=8.4 dB ±1.6).

The 18 vocalizations and 4 backgrounds (no context, noise, clean guitar, distortion guitar) were then superimposed to create 72 different stimuli (onset of the vocalization = 30 ms. after the onset of the context, See Figure 1- bottom). While we were primarily interested in context effects, the rationale for varying both cues to vocal arousal and stimulus contexts was to avoid experimental demand effects induced by situations where context is the only manipulated variable.

### Procedure

On each trial, participants listened to one stimulus (vocalization + background) and evaluated its perceived emotional arousal and valence on two continuous sliders positioned below their respective self-assessment manikin (SAM) scales (Bradley & Lang, 1994): a series of pictures varying in both affective valence and intensity, that serve as a nonverbal pictorial assessment technique directly measuring a person’s affective reaction to a stimulus. Participants were instructed to evaluate the emotional state of the speaker, ignoring background (noise or guitar). A first training block was presented, with 20 trials composed of vocalizations with no background (randomly selected from the subsequent set of stimuli). After this phase, listeners received a score out of 100 (actually a random number between 70 and 90) and were asked to maintain this performance in subsequent trials, despite the sounds being thereafter embedded in background noise. Each of the 18 vocalizations (2 speakers x 3 pitches x 3 anger levels) was then presented five times in four different contexts (none, clean guitar, distortion guitar, noise), with a 1s inter-trial interval. The main experiment included 360 randomized trials divided into 6 blocks. To motivate continued effort, participants were asked to maximize accuracy during the practice phase, and at the end of each block they received fake feedback scores, on average slightly below that of the training phase (a random number between 60 and 80). Participants were informed that they could receive a financial bonus if their accuracy was above a certain threshold (all participants received the bonus regardless of their score). The experiment was run in a single session lasting 75 minutes. Sound files were presented through headphones (Beyerdynamic DT 270 PRO, 80 ohms), with a fixed sound-level that was judged to be comfortable by all the participants.

### Statistical analyses

We used the continuous slider values corresponding to the two SAM scales (coded from 0 to 100 between their two extremes) to extract valence and emotional arousal ratings. We analyzed the effect of context on these ratings with two repeated-measures ANOVAs, conducted separately on negativity (100-valence) and emotional arousal, using level of portrayed vocal arousal in the voice (3 levels) and context (4 levels: Voice-alone, clean-guitar, distorted guitar, noise) as independent variables. Post-hoc tests were Tukey HSD. All statistical analyses were conducted using R (R Core Team, 2013). Huynh-Feldt corrections 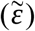 for degrees of freedom were used where appropriate. Effect sizes are reported using partial eta-squared *η*_*p*_^*2*^.

## 2. Results

To control that stimuli with increasing levels of portrayed vocal arousal indeed had more nonlinearities, we subjected the 18 vocal stimuli to acoustic analysis with the Praat software (Boersma, 2011), using three measures of voice quality (jitter, shimmer, noise-harmonic ratio) commonly associated with auditory roughness and noise. All three measures scaled as predicted with portrayed vocal arousal (Figure 1-top).

Repeated-measures ANOVAs revealed a main effect of portrayed vocal arousal on judgments of emotional valence, 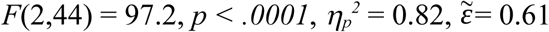, with greater levels of speaker anger associated with more negative valence; a main effect of context 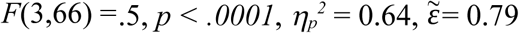, with sounds presented in distorted guitar judged more negative than in clean guitar, and sounds presented in noise judged more negative than in distorted guitar (Figure 2-top, left). Ratings of stimuli presented alone or in distorted guitar did not differ (Tukey HSD). Context did not interact with the effect of anger on perceived valence (Figure 2-top, right).

**Figure 2:**
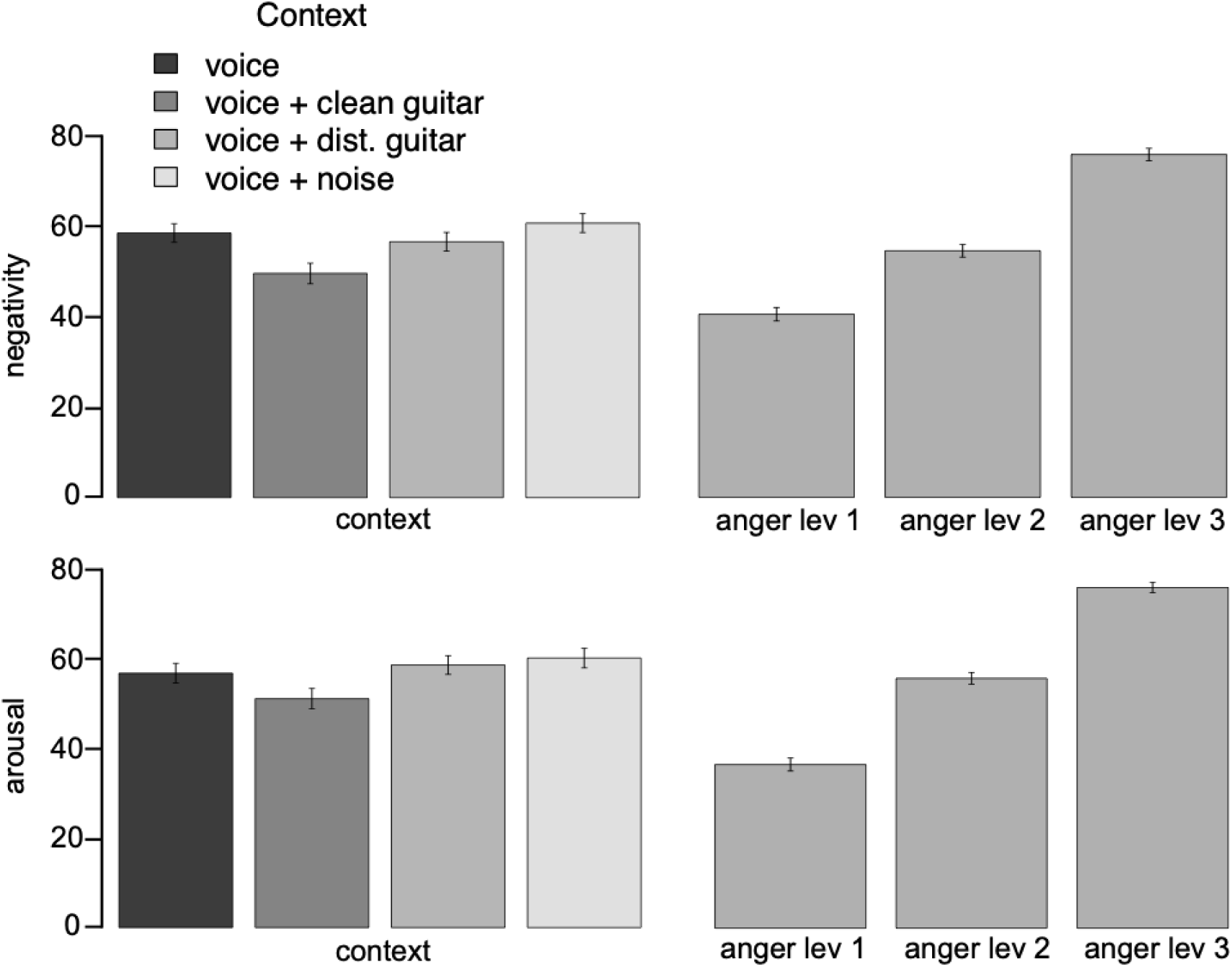
Perceived valence (negativity) and emotional arousal as a function of three levels of portrayed vocal arousal of human vocalizations presented in different sound contexts. Error-bars show 95% confidence intervals on the mean.

A similar pattern of results was found for judgments of emotional arousal, with a main effect of portrayed vocal arousal, 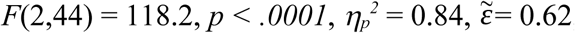, increasing perceived emotional arousal, and a main effect of context 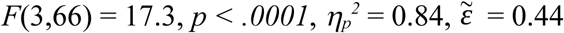. Sounds presented with noise and distorted guitar were judged as more emotionally aroused than with clean guitar (Figure 2-bottom, left). Ratings in noise and distorted guitar contexts did not differ (Tukey HSD) and, as above, context did not interact with the effect of anger on perceived emotional arousal (Figure 2-bottom, right).

## 3. Discussion

As expected, voices with higher levels of portrayed anger were judged as more negative and more emotionally aroused than the same voices produced with less vocal arousal. Both the perceived valence and emotional arousal of voices with high vocal arousal were significantly affected by both musical and non-musical contexts. However, contrary to what would be predicted e.g. by the aesthetic enjoyment of nonlinear vocal sounds in rough musical textures by death metal fans (Thompson, Geeves & Olsen, 2018) or more generally, the suggestion of the top-down deactivation of the effect of nonlinear features in appropriate musical contexts (Zatorre & Belin, 2015), we did not find any incongruency effect between musical contexts and vocal signals, and in particular did not find support for the notion that the presence of a distorted guitar would reduce emotional effects of nonlinearities in voices. Instead, screams with high levels of portrayed vocal arousal were perceived as more negative and more emotionally aroused in the context of background nonlinearities. This suggests that local judgments such as the emotional appraisal of one isolated part of an auditory scene (e.g., a voice) are computed on the basis of the global acoustic features of the scene. It is possible that this effect results from a high-level integration of cues similar to the emotional evaluation of audiovisual signals (de Gelder & Vroomen, 2000), in which both signal and context are treated as coherent communicative signals reinforcing each other. Another possibility is that stream segregation processes cause the nonlinear features of the vocal stream to perceptively fuse with that of the other (i.e., musical or noise) stream (see e.g. Vannier et al., 2018).

These results raise several questions for future research. First, to understand the neural time-course of these contextual effects as well as disentangle the respective contributions of subcortical (e.g. amygdala; Arnal et al., 2015) and high-level decision processes, the current study could be complemented with fMRI imaging and MEG analysis. Second, future research could explore psychoacoustical sensitivity to vocal roughness, and how discrimination thresholds might shift as a function of low-level properties of the acoustic context. Additionally, error management principles could be at play as listeners might be biased to over-detect nonlinear-based roughness in cases of possible danger (Blumstein, Whitaker, Kennen, & Bryant, 2017). Finally, research could examine whether context effects exist with signals that are not conspecific vocalizations, such as other animals’ vocalizations/calls or synthetic alarm signals.

More generally, these findings suggest that musical features are processed similarly to vocal sounds, and that affective information in music is treated as a communicative signal (Bryant, 2013). Instrumental music constitutes a recent cultural innovation that often explicitly imitates properties of the voice (Juslin & Laukka, 2003). The cultural evolution of music technology and sound generation exploits perceptual mechanisms designed to process communicative information in voices. These results provide an excellent example of how an ecologically-based theoretical framework can be used to understand what might otherwise appear to be novel features in contemporary cultural phenomena such as vocal and instrumental music.

## Acknowledgements

The authors thank Hugo Trad for help running the experiment. All data collected at the Sorbonne-Université INSEAD Center for Behavioural Sciences.

## Author contributions

ML and EP made equal contributions for conceiving the study, recording and designing stimuli, running the experiment, analyzing data, and drafting the manuscript. GB contributed to the interpretation of the data analysis and formalization of the results. JJA contributed to the conception of the experimental design. All authors contributed to the writing of the manuscript.

## Data accessibility

Matlab files and stimuli to run the experiment, R files and python notebook to analyze the results, as well as a .fxb file for the guitar distortion plugin are available at the following URL: https://nubo.ircam.fr/index.php/s/QAMG78HPymso26o

